# Cysteine signaling in plant pathogen response

**DOI:** 10.1101/2025.03.13.642987

**Authors:** Jannis Moormann, Björn Heinemann, Cecile Angermann, Anna Koprivova, Ute Armbruster, Stanislav Kopriva, Tatjana M. Hildebrandt

**Affiliations:** Institute for Plant Sciences, Cluster of Excellence on Plant Sciences (CEPLAS), University of Cologne, Zülpicher Straße 47a, 50674 Cologne, Germany; Molecular Photosynthesis, Cluster of Excellence on Plant Sciences (CEPLAS), Heinrich-Heine-University Düsseldorf, Universitätsstraße 1, D-40225, Düsseldorf, Germany

**Keywords:** amino acid metabolism, sulfur signaling, infochemicals, immunometabolism, mitochondria, proteomics

## Abstract

The amino acid cysteine is the precursor for a wide range of sulfur-containing functional molecules in plants, including enzyme cofactors and defense compounds. Due to its redox active thiol group cysteine is highly reactive. Synthesis and degradation pathways are present in several subcellular compartments to adjust the intracellular cysteine concentration. However, stress conditions can lead to a transient increase in local cysteine levels. Here we investigate links between cysteine homeostasis and metabolic signaling in *Arabidopsis thaliana*. The systemic proteome response to cysteine feeding strongly suggests that Arabidopsis seedlings interpret accumulation of cysteine above a certain threshold as a signal for a biotic threat. Cysteine supplementation of Arabidopsis plants via the roots increases their resistance to the hemibiotrophic bacterium *Pseudomonas syringae* confirming the protective function of the cysteine induced defense pathways. Analysis of mutant plants reveals that the balance of cysteine synthesis between the cytosol and organelles is crucial during Arabidopsis immune response to *Pseudomonas syringae*. The induction profile of pathogen responsive proteins by cysteine provides insight into potential modes of action. Our results highlight the role of cysteine as a metabolic signal in the plant immune response and add evidence to the emerging concept of intracellular organelles as important players in plant stress signaling.

## Introduction

Amino acids have multiple functions in plants. In addition to their role during protein biosynthesis, they are an integral part of several biosynthetic pathways and involved in signaling processes including plant stress responses (Heinemann & Hildebrandt, 2021). Amino acids can serve as markers reflecting nutrient availability or as carbon source for alternative respiratory pathways under energy starvation (Pedrotti et al., 2018). Proline is known to function as an osmolyte during osmotic stress and as molecular chaperone preventing protein aggregation (Szabados & Savouré, 2010). In addition, amino acids serve as precursors for various molecules involved in plant immunity. Perturbations in amino acid metabolism have been reported to affect plant immune responses in various ways (Moormann et al., 2022). The phloem-mobile signaling molecule N-hydroxypipecolic acid, required for establishing systemic acquired resistance, is synthesized from lysine (R. N. Gupta & Spenser, 1969; Návarová et al., 2012). Aromatic amino acids are precursors for a broad range of specialized molecules that crucially shape the interaction between plants and microbes such as coumarins, phytoalexins and indolic glucosinolates (Glawischnig, 2007; Harun et al., 2020; Maeda & Dudareva, 2012; Pastorczyk et al., 2020). The thiazole ring of camalexin, the characteristic phytoalexin of *Arabidopsis thaliana*, originates from the cysteine residue of glutathione (Su et al., 2011).

Among amino acids, cysteine is unique as it occupies a central position in plant sulfur metabolism. It is the precursor for a wide range of sulfur-containing molecules in the cell, including methionine, essential vitamins and cofactors such as thiamin, lipoic acid, biotin, Fe-S clusters and molybdenum cofactor (Droux, 2004; Giovanelli et al., 1985; van Hoewyk et al., 2008). Furthermore, the tripeptide γ-glutamyl-cysteinyl-glycine, also known as glutathione, relies on the incorporation of cysteine to function as the main determinant of cellular redox homeostasis (Foyer & Noctor, 2011). Its antioxidant property is based on the redox potential of the thiol group. Next to oxidative stress protection, glutathione also acts in detoxification of heavy metals and xenobiotics as well as in the plant defense response (Noctor et al., 2024). In proteins, cysteine residues contribute to structure, stability and function. When located in the active sites of enzymes they are essential for catalysis of many enzymatic reactions as well as metal cofactor binding. Moreover, thiol groups can undergo oxidation to form covalent disulfide bridges, aiding protein folding and stability. Reversible oxidation/reduction of these disulfide bridges poses a mechanism for redox regulation of proteins (Buchanan & Balmer, 2005). Another layer of regulation is added by posttranslational cysteine modifications such as glutathionylation, nitrosylation, sulfenylation and persulfidation (Begara-Morales et al., 2016; Moseler et al., 2024).

Cysteine is the product of the plant sulfur assimilatory pathway. Sulfate is taken up from the soil by specific transporters and activated by ATP sulfurylase. The resulting adenosine-5‘-phosphosulfate (APS) is reduced in a two-step reaction via sulfite to sulfide by APS reductase and sulfite reductase. Sulfide is incorporated into O-acetylserine (OAS) by OAS-(thiol)lyase (OASTL) to produce cysteine (Takahashi, 2010; Takahashi et al., 2011). The amino acid precursor OAS is synthesized by serine acetyltransferase (SERAT) from serine and acetyl-CoA. OASTL and SERAT form a cysteine synthase complex which is required for regulating cysteine synthesis based on substrate availability (Droux, 2003; Wirtz & Hell, 2006). Both enzymes have isoforms localized in the cytosol (SERAT1;1, SERAT3;1, SERAT3;2, OASTL-A), plastids (SERAT2;1, OASTL-B) and mitochondria (SEART2;2, OASTL-C) enabling OAS and cysteine synthesis in different subcellular locations (Hell & Wirtz, 2011; Ruffet et al., 1995; Watanabe et al., 2008). Cysteine desulfurases, which transfer sulfur to Fe-S cluster scaffold proteins, are present in plastids and mitochondria as well (Couturier et al., 2013). The cytosolic cysteine desulfurase ABA3 is required for molybdenum cofactor synthesis (Caubrière et al., 2023). Cysteine levels are tightly regulated and generally kept low within the cell requiring adequate rates of cysteine degradation (Hildebrandt et al., 2015). In the cytosol, the L-cysteine desulfhydrase DES1 deaminates cysteine resulting in pyruvate, ammonia and sulfide (Alvarez et al., 2010). In mitochondria, a four-step process catalyzing the complete oxidation of cysteine to pyruvate and thiosulfate is facilitated by an unknown aminotransferase, the 3-mercaptopyruvate sulfur transferase STR1, and the sulfur dioxygenase ETHE1 (Höfler et al., 2016). Cysteine levels vary among cellular compartments and are highest in the cytosol which is known to be the main contributor of cysteine with concentrations around 300 µM (Heeg et al., 2008; Krüger et al., 2009). Accordingly, OASTL-A together with OASTL-B were found to account for 95% of OASTL activity in *Arabidopsis* protein extracts, while OASTL-C contributed only 5% of the activity. However, all three major OASTL isoforms were shown to largely compensate for each other’s absence in null mutant studies (Birke et al., 2013; Heeg et al., 2008).

Previous findings provide some insight into the relevance of compartment specific cysteine metabolic pathways. Cytochrome c oxidase, the last enzyme of the mitochondrial respiratory chain, is strongly inhibited by cyanide as well as by hydrogen sulfide (Cooper & Brown, 2008). Biosynthesis of camalexin and the gaseous phytohormone ethylene both lead to the production of cyanide in non-cyanogenic plants such as *Arabidopsis* (Böttcher et al., 2009; Peiser et al., 1984). Cyanide detoxification in the mitochondria is achieved by conversion of cysteine and cyanide to hydrogen sulfide and β-cyanoalanine catalyzed by cyanoalanine synthase (CAS-C1) (Hatzfeld et al., 2000; Watanabe et al., 2008). Hydrogen sulfide in turn is detoxified via incorporation into cysteine by mitochondrial OASTL-C, creating a cyclic detoxification pathway. OASTL-C might also be involved the regulation of sulfur homeostasis since loss or decreased activity due to a single nucleotide polymorphism in *Arabidopsis* accessions leads to reduced sulfate uptake (Koprivova et al., 2023). In addition, cysteine synthesis was found to play a role during stomatal closure in response to drought stress. Upon soil drying, sulfate is transported to the guard cells via the xylem and is incorporated into cysteine (Ernst et al., 2010). Subsequently, cysteine mediates abscisic acid (ABA)-dependent stomatal closure in multiple ways (Heinemann & Hildebrandt, 2021). It is a substrate of the MoCo-sulfurylase ABA3 involved in ABA synthesis (Caubrière et al., 2023). ABA induces the expression of DES1, which uses cysteine as a substrate to produce hydrogen sulfide in the cytosol (Chen et al., 2020). Hydrogen sulfide accumulation leads to persulfidation and activation of several proteins involved in ABA-induced stomatal closure including kinases and transcription factors (Chen et al., 2020; Shen et al., 2020; M. Zhou et al., 2021). These findings illustrate the wide range of functions and different modes of action of cysteine and its related metabolism. Taken together, the versatile chemical nature of cysteine and the high degree of compartmentalization of its metabolism further highlight its potential for multiple, yet unknown, regulatory functions.

In order to identify additional potential signaling functions of cysteine in *Arabidopsis*, we analyzed the response of seedlings to an artificial increase in cysteine levels. The proteome signature of the cysteine treated seedlings indicated perception of the disturbance in cysteine homeostasis as a biotic threat. Thus, we further investigated the role of compartment specific cysteine metabolism in the interaction of *Arabidopsis* plants with the leaf pathogen *Pseudomonas syringae* pv tomato DC3000 (*Pst*) and identified cytosolic and mitochondrial cysteine synthesis as major contributors to pathogen resistance.

## Results

### The seedling proteome response to increased cysteine concentrations indicates biotic stress signaling

To understand the systemic response of *Arabidopsis thaliana* to an increased cysteine content we performed a feeding experiment. Six-day-old seedlings grown in liquid culture were supplemented with 1 mM L-cysteine for 24h (Fig. 1A, Suppl. Fig. S1A). This treatment led to a 7.6-fold increase in the seedling cysteine content, and the glutathione content was also significantly higher (1.7 -fold) than in control seedlings (Fig. 1B). The cysteine content in the medium continuously decreased most likely due to a combination of uptake by the plants and chemical oxidation processes in the medium and was completely depleted at the end of the 24h incubation time (Suppl. Fig. S1B). To test, whether cysteine oxidation led to a depletion of oxygen in the medium or had an effect on seedling respiration we analyzed the oxygen content of the medium as well as seedling oxygen consumption rates, but did not detect any significant differences between cysteine treatment and control (Suppl. Fig. S1C).

**Fig. 1:**
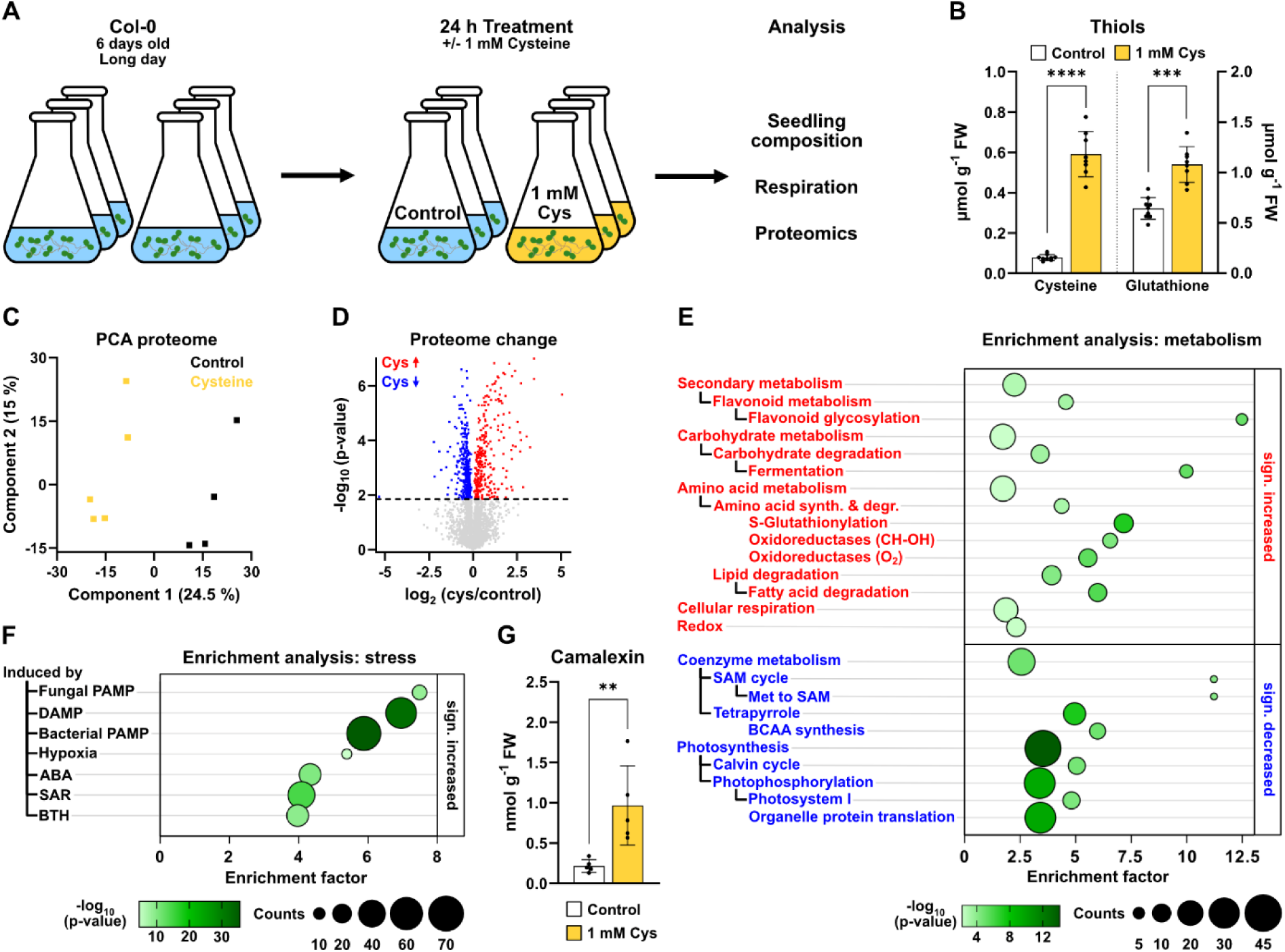
**Seedling proteome response to cysteine treatment**. (**A**) An *A. thaliana* seedling culture was supplemented with 1 mM L-cysteine for 24 hours before harvest. (**B**) Seedling content of L-cysteine and reduced glutathione (GSH) [µmol · g fresh weight ^-1^] (n = 8-9). (**C**) Principal component analysis of shotgun proteomics dataset with control (black) and cysteine-treated (yellow) seedlings. (**D**) Volcano plot illustrating differences in the proteome of cysteine treated seedlings vs. controls. Proteins of significantly higher or lower abundance after cysteine feeding are marked in red and blue, respectively (t-test, FDR < 0.05). (**E**) Enrichment of functional categories in proteins that are significantly increased (top, red) or decreased (bottom, blue) in the presence of cysteine. (**F**) Enrichment of stress induced categories among proteins that are significantly increased in the presence of cysteine. ABA, abscisic acid; BCAA, branched-chain amino acid; BTH, benzothiadiazole, DAMP, damage-associated molecular pattern; JA, jasmonic acid; PAMP, pathogen-associated molecular pattern; SAM, S-adenosylmethionine; SAR, systemic acquired resistance. The complete proteomics dataset including the enrichment analysis is provided as Suppl. Dataset S1. (**G**) Seedling camalexin content [nmol · g fresh weight-1] (n = 5). Mean (bars) and individual (dots) values ± SD are shown. Asterisks indicate statistically significant differences compared with control seedlings following students t-test (** P < 0.01 > 0.001; *** P < 0,001 > 0,0001; **** P < 0,0001). Raw data is provided in Suppl. Dataset S4.

Shotgun proteome analysis revealed a distinct effect of cysteine feeding on the composition of the seedling proteome (Fig. 1C). 479 of the 4613 detected protein groups were significantly increased and 670 significantly decreased in cysteine treated compared to control samples (Fig. 1D). Among the strongly downregulated proteins were the transcriptional activator Hem1, kinases associated with ABA signaling, and proteins involved in sulfur assimilation whereas glutathione-S transferases (GSTs) as well as UDP-glucosyltransferases were most drastically increased (Suppl. Dataset S1). In order to systematically identify major features of the proteome response to increased cysteine levels, we performed an enrichment analysis on functional annotations of the significantly changed proteins (Suppl. Dataset S1). Enrichment of metabolic pathways was analyzed based on a modified version of the MapMan annotation system (https://mapman.gabipd.org; Fig. 1E). The results indicated a decrease in pathways related to photosynthesis including photophosphorylation, the Calvin-Benson-Bassham cycle and also tetrapyrrole and organellar protein synthesis (Fig. 1E, blue). In contrast, pathways required for heterotrophic energy metabolism such as carbohydrate and lipid degradation as well as cellular respiration showed increased abundance after cysteine treatment (Fig. 1E, red). The enrichment of stress-related categories (glucosyltransferases, oxidoreductases, glutathione-S-transferases) among the significantly increased proteins prompted us to systematically probe our dataset for characteristic stress response profiles. To this end, we performed an additional enrichment analysis on the basis of published transcriptome profiles for diverse abiotic and biotic stress conditions as well as elicitor and hormone treatments focusing on the proteins significantly increased after cysteine feeding (Fig. 1F, Suppl. Dataset S1). Strikingly, six of the seven significantly enriched categories in cysteine treated seedlings were associated with pathogen response. Enriched categories of proteins were those induced by pathogen-associated molecular patterns (PAMPs), damage-associated molecular patterns (DAMPs), systemic acquired resistance (SAR), treatment with jasmonic acid (JA) or the salicylic acid analogon benzothiadiazole (BTH) strongly suggesting a role for cysteine during biotic stress signaling (Fig. 1F). Indeed, the concentration of the phytoalexin camalexin was significantly (4.5-fold) increased in the cysteine-treated seedlings indicating a similarity to an active pathogen response (Fig. 1G).

### Cysteine treatment induces pathogen resistance

To test the physiological relevance of the observed induction of pathogen response pathways by cysteine feeding under physiological conditions we watered six-week-old *A. thaliana* plants grown on soil with 10 mM cysteine 24 hours before performing a pathogen assay using the hemibiotrophic bacterium *Pseudomonas syringae* pv. tomato DC3000 (*Pst*) (Fig. 2A). Quantification of thiols confirmed that cysteine was taken up by the plants and transported to the leaves (Fig. 2B). Cysteine accumulated in the rosette leaves to a similar extent as in the seedling culture in treated compared to control plants (7-fold). The cysteine treated plants were significantly less susceptible to *Pst* indicated by reduced bacterial growth and less chlorosis after three days of infection (Fig. 2C,D).

**Fig. 2:**
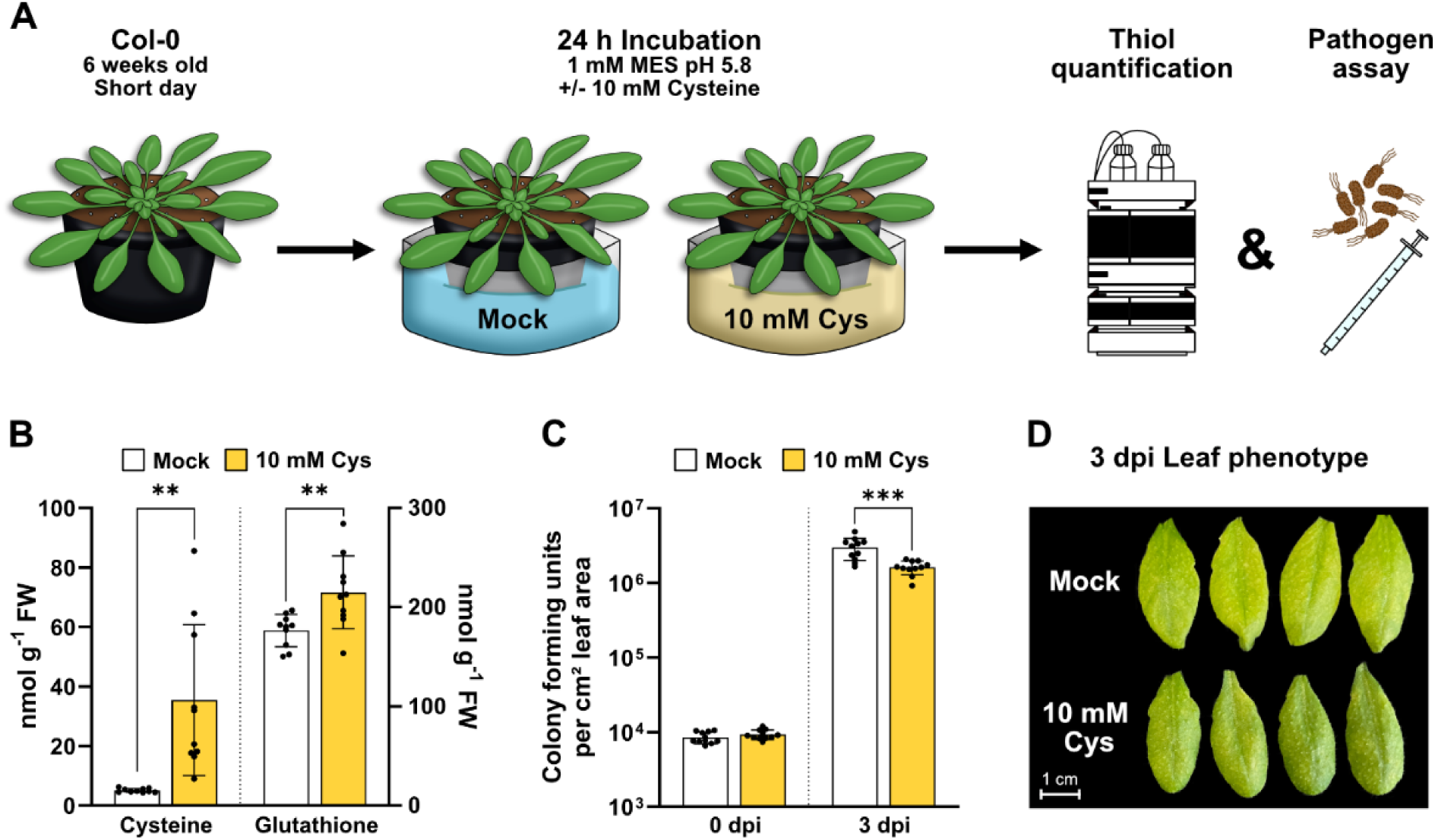
**Cysteine treatment induces resistance to the virulent pathogen *Pseudomonas syringae***. (**A**) Schematic representation of the workflow: *A. thaliana* plants were grown for 6 weeks under short-day conditions and incubated in 10 mM L-cysteine for 24 h before further analysis. (**B**) Content of L-cysteine and reduced glutathione (GSH) [nmol · g fresh weight^-1^] in the rosette leaves of mock treated (white bars) and cysteine treated (yellow bars) plants (n = 10) (**C**) *Pst* infection assay: Four leaves per plant were infiltrated with *P. syringae* DC3000 solution including 5 × 10^5^ colony forming units (CFU) per ml using a needleless syringe and sampled at 0 and 3 days past infection (dpi) to quantify CFU per cm leaf area (n = 11). (**D**) Phenotype of four representative leaves per treatment at three days after inoculation. Mean (bars) and individual (dots) values ± SD are shown. Asterisks indicate statistically significant differences compared with mock-treated plants (** P < 0.01 > 0.001; *** P < 0,001 > 0,0001). Raw data is provided in Suppl. Dataset S4.

### Pathogen attack induces cysteine synthesis, and proteome responses to cysteine treatment and pathogen interaction overlap

To investigate the function of cysteine during plant-pathogen interaction, we quantified thiols in *Arabidopsis* leaves infected with *Pst*. Cysteine levels were significantly increased already 12 hours post infection (hpi), peaked at 24 hpi at 6.6-fold level of the mock treated controls and remained constantly high until 48 hpi (Fig. 3A). The proteome of the infected leaves showed clear differences to the mock treated samples (Fig. 3B; Suppl. Dataset S2). The pattern of changes in individual protein abundances was highly consistent between the two timepoints analyzed with a stronger response at 48 hpi (Suppl. Dataset S3; Suppl. Fig. S2). 697 proteins were significantly increased and 582 proteins significantly decreased at both timepoints. Highlighting proteins with significant changes in the cysteine-treated seedlings in the *Pst* infection dataset illustrates that there is also a strong overlap between the *Arabidopsis* proteome response to increased cysteine contents and pathogen attack (Fig. 3C; Suppl. Fig. S2; Suppl. Dataset S3). 174 proteins were significantly increased and 268 proteins significantly decreased after cysteine feeding as well as after pathogen attack. Among the consistently highly induced proteins in both responses were several glutathione-S-transferases, UDP-glucosyltransferases, and ALTERNATIVE OXIDASE 1A (Suppl. Dataset S3). Enrichment analysis of functional annotations identifies a repression of photosynthesis related pathways, coenzyme metabolism as well as organellar protein translation (Fig. 3D, blue) and an induction of lipid catabolism (Fig. 3D, red) as common effects of both treatments.

**Fig. 3:**
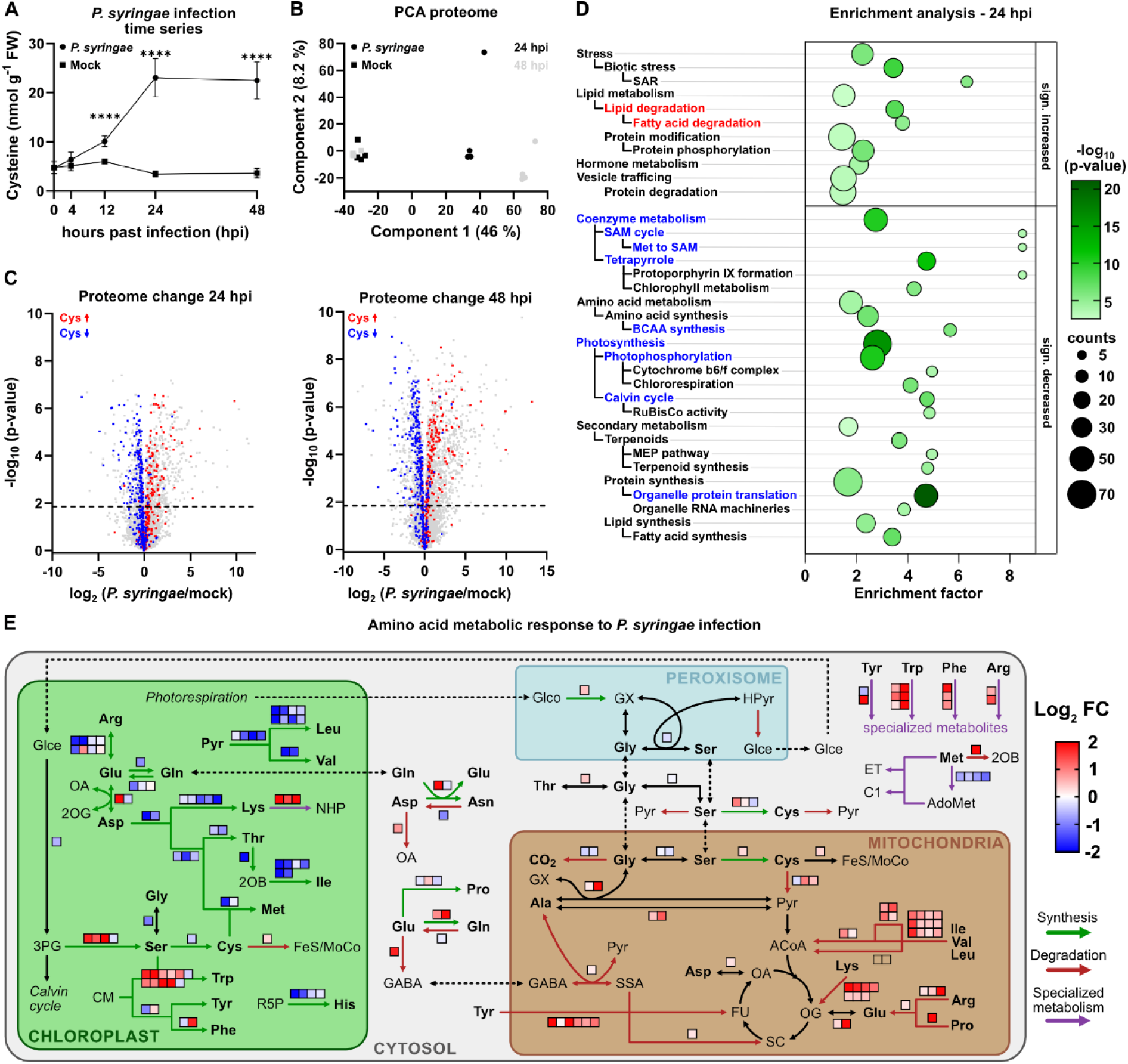
**Cysteine accumulation and proteome response during pathogen interaction**. Six weeks old *A. thaliana* plants were infiltrated with *P. syringae* DC3000 (*Pst*) (2.5 × 10^6^ colony forming units per ml). Six leaves per plant were sampled at the indicated time points. (**A**) Content of L-cysteine [nmol · g fresh weight^-1^] in infected vs. mock treated leaves (n = 4-9). Mean values ± SD are shown. Asterisks indicate statistically significant differences compared with mock-treated plants (**** P < 0,0001). (**B**) Principal component analysis of the proteome of *Pst*- and mock-infected leaves (circles and squares, respectively) at 24 and 48 h after inoculation (black and grey, respectively) (**C**) Volcano plots illustrating differences in the proteome of *Pst*- vs. mock-infected plants at 24 and 48 hpi. Proteins of significantly higher or lower abundance after treatment with 1 mM L-cysteine (Fig. 1C) are highlighted in red and blue, respectively. (**D**) Enrichment of functional categories in proteins that are significantly increased (top) or decreased (bottom) at 24 hpi with *Pst*. Red and blue fonts indicate significantly increased or decreased categories, respectively, that were also enriched after cysteine treatment (Fig. 1E). (**E**) Effect of *Pst* infection on amino acid metabolic pathways. Relative protein abundances in infected vs. mock treated leaves at 48 hours after inoculation *Pst.* Proteins significantly increasing or decreasing in abundance are indicated by red or blue squares, respectively. 2-OB, 2-oxobutyrate; 2-OG, 2- oxoglutarate; 3PG, 3-phosphoglycerate; ACoA, Acetyl-CoA; AdoMet, S-Adenosylmethionin; C1, C1-metabolism; CM, chorismate; ET, ethylene; FeS, iron-sulfur cluster; FU, fumarate; GABA, γ-aminobutyric acid; Glce, glycerate; Glco, glycolate; GX, glyoxylate; Hpyr, Hydroxypyruvate; MoCo, molybdenum cofactor; NHP, N-hydroxypipecolic acid; OA, oxaloacetic acid; Pyr, pyruvate; R5P, ribose-5-phosphate; SSA, succinic semialdehyde. The complete proteomics dataset including the enrichment analysis is provided in Suppl. Dataset S2. Additional raw data is provided in Suppl. Dataset S4.

We performed additional experiments to validate these cysteine responses with potential relevance in immune signaling (Suppl. Figure S3). The chlorophyll content of the cysteine treated seedlings was significantly decreased by 10% compared to the control, which is in line with the repression of photosynthesis indicated by the proteome data (Suppl. Figure S3A). However, we did not detect any significant effect of cysteine supplementation via the roots on the photosynthetic performance of the leaves as determined by chlorophyll a fluorescence analysis (Suppl. Figure S3B). The overall seedling composition with respect to carbohydrates, lipids and proteins remained unchanged (Suppl. Figure S3C). The induction of several glutathione-S-transferases (GSTs) in response to cysteine treatment led to a 1.6fold increase in total GST abundance, which was also reflected in a 1.7fold higher GST activity (Suppl. Figure S3D). To evaluate, whether the trigger for the induction of GSTs and other stress related proteins such as alternative oxidase might be an accumulation of reactive oxygen species we performed DAB staining with the seedlings (Suppl. Figure S3E). The results revealed that hydrogen peroxide levels in the cysteine treated seedlings were even lower than in the controls indicating that other signaling processes are involved.

Several aspects of the proteome response to *Pst* interaction such as a decrease in terpenoid synthesis and an increase in hormone metabolism or vesicle trafficking did not become apparent during cysteine feeding and thus might be unrelated to cysteine signaling. The general effect on amino acid metabolism included an increase in the catabolic pathways mainly localized in the mitochondria and a decrease in plastidic synthesis pathways. A clear exception were the synthesis pathway of serine and the aromatic amino acids, which were strongly induced together with downstream pathways required for their conversion to specialized metabolites (Fig. 3E).

### Compartment specific cysteine synthesis is required for pathogen resistance

A focus on the proteins involved in cysteine metabolism illustrates that the cytosolic and the mitochondrial pathway for cysteine synthesis as well as plastidic GSH production increased in abundance after pathogen attack whereas sulfate reduction and cysteine synthesis in the chloroplasts decreased (Fig. 4A). In order to estimate the contributions of the individual compartments to cellular cysteine production we compared the total abundance of O-acetyl serine lyase (OASTL) isoforms catalyzing the last step in cysteine synthesis in the leaf proteome (Fig. 4B). The cytosolic and plastidic isoforms OASTL-A and OASTL-B were of similar abundance, but OASTL-C localized in the mitochondria represented only about 5% of the total OASTL leaf content. During pathogen interaction OASTL-A and OASTL-C significantly increased in abundance whereas OASTL-B significantly decreased (Fig. 4A). Next, we tested the response of knockout mutant lines for the individual OASTL isoforms during interaction with *Pst* (Fig. 4C-G). The *oastl-c* (mitochondrial) mutant was significantly more susceptible to *Pst* infection compared to the wild type and the oastl-b (chloroplast) mutant (Fig. 4C). The stronger infection was also visible in the phenotype of the infected leaves, which were shriveled up with necrotic tips (Fig. 4D). The cytosolic *oastl-a* mutant showed an intermediate level of bacterial growth rates between the wild type and *oastl-c* (Fig. 4C). We detected a significant decrease in leaf camalexin levels at 24 hpi in *oastl-a* and *b* and the same tendency in *oastl-c*. (Fig. 4E). Total leaf cysteine levels were lower than in the wild type already under control conditions in *oastl-a* (deficient in cytosolic cysteine synthesis) and also the cysteine increase during infection lagged behind in this line but not in the others (Fig. 4F, grey bars). Glutathione levels showed a similar trend (Fig. 4G).

**Fig. 4:**
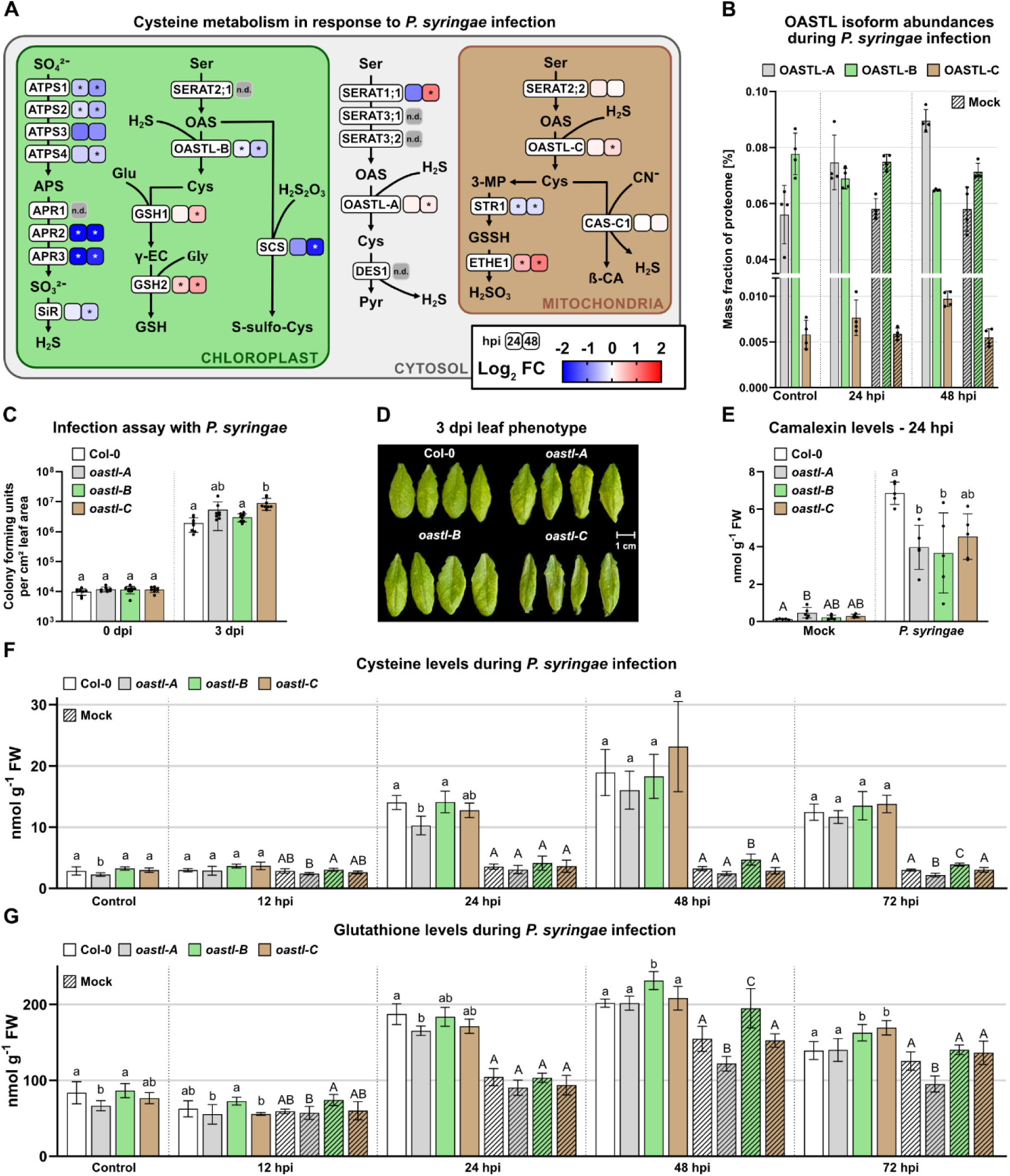
**Compartmentalization of cysteine synthesis in pathogen resistance**. (**A**) Effect of *Pst* infection on cysteine metabolic pathways: Relative protein abundances in infected vs. mock treated leaves at 24 and 48 hours after inoculation with *P. syringae* DC3000 (*Pst*) (2.5 × 10^6^ colony forming units (CFU) per ml). Proteins increasing or decreasing in abundance are indicated by red or blue squares, respectively (n.d., undetected proteins). Asterisks indicate statistically significant differences compared with mock treated plants following students t-test (*P < 0.05). APS, adenosine 5‘-phophosulfate; ATPS, ATP sulfhydrase; APR, APS reductase; SiR, sulfite reductase; SERAT, serine acetyltransferase; OAS, O-acetylserine; OAS-A, O-acetylserin(thiol)lyase; GSH, glutathione; GSH1, γ-glutamyl-cysteine synthetase; GSH2, GSH syntethase; SCS, S-sulfocysteine synthase; DES1, L-cysteine desulfhydrase; Pyr, Pyruvate; 3-MP, 3-mercaptopyruvate; STR1, 3-MP sulfurtransferase; GSSH, GSH-persulfide; ETHE1, sulfur dioxygenase; β-CA, β-cyanoalanine; CAS-C1, CAS synthase. (**B**) Total abundance of O-acetylserin(thiol)lyase isoforms in control, mock treated and *Pst* infected leaves of wild type *A. thaliana* plants calculated from quantitative iBAQ-values (n = 4). (**C**) *Pst* infection assay: Four leaves per plant were infiltrated with *P. syringae* DC3000 solution including 5 × 10^5^ colony forming units (CFU) per ml using a needleless syringe and sampled at 0 and 3 days past infection (dpi) to quantify CFU per cm leaf area. (n = 8). (**D**) Phenotype of four representative leaves per treatment at three days after inoculation. (**E**) Camalexin content [nmol · g fresh weight^-1^] in mock treated and *Pst* infected leaves of wild type *A. thaliana* plants and OASTL-deficient mutant lines (n = 4-5). (**F**) Cysteine and (**G**) Glutathione content [nmol · g fresh weight^-1^] in control, mock treated, and *Pst* infected leaves of wild type *A. thaliana* plants and OASTL-deficient mutant lines (n = 5-10). Mean (bars) and individual (dots) values ± SD are shown. Letters indicate statistically significant differences following ANOVA with Tukey’s test (α = 0.05). Raw data is provided in Suppl. Dataset S4.

## Discussion

### Links of cysteine homeostasis to plant immune signaling

The systemic proteome response to cysteine treatment strongly suggests that *Arabidopsis* seedlings interpret accumulation of cysteine above a certain threshold as a signal for a biotic threat. In accordance with these results, artificially increasing the intracellular cysteine concentration in mature plants renders them significantly less susceptible to infection by the hemibiotrophic bacterium *Pseudomonas syringae,* confirming the protective function of the defense pathways activated in response to cysteine treatment. Cysteine strongly accumulated within the first 24h of infection in control plants that had not been treated prior to inoculation showing that a disturbance in cysteine homeostasis is an integral feature of the interaction between plant and pathogen. These findings indicate that the results obtained using the artificial but also highly reproducible and homogenous seedling culture approach can provide valuable insight into physiologically relevant processes. Effects observed consistently in seedling cultures and in the plants grown on soil despite the differences in age, tissues, and cultivation style have the potential to reveal fundamental aspects of metabolic immune signaling.

Several proteins were strongly induced by both, cysteine feeding as well as infection with *Pst* and thus might provide some insight into potential links between cysteine homeostasis and immune signaling. The functional category with the strongest enrichment in cysteine induced proteins consisted of UDP-glucosyltransferases (UGTs) of the 73B subfamily annotated as flavonoid glycosidases. Several additional UGTs responded similarly to *Pst* infection and cysteine treatment, and among them UGT71C5, UGT74F2, and UGT87A2 showed the strongest consistent induction (Suppl. Dataset S3). The multigene family of uridine diphosphate-dependent glucosyltransferases catalyzes the covalent addition of sugars from nucleotide UDP sugar donors to functional groups on a variety of compounds which can increase the solubility of lipophilic metabolites or inactivate hormones and signaling molecules (Lairson et al., 2008). A number of UGTs have already been linked to plant immunity. UGT74F2 and UGT76B1 glycosylate and thereby inactivate immune signals including salicylic acid (SA) and N-hydroxypipecolic acid. Knockout lines develop an autoimmune phenotype indicating that this function is required for balancing and containing immune responses (Bauer et al., 2021; Lim et al., 2002). UGT71C5 uses the phytohormone abscisic acid (ABA) as a substrate and is involved in regulating ABA homeostasis (Liu et al., 2015). ABA mediates abiotic stress responses and acts antagonistically to SA induced immune signaling (Torres Zabala et al., 2009; Torres-Zabala et al., 2007). The induction of UGT71C5 by cysteine will therefore amplify the plant pathogen response by suppressing antagonistic signals. A modeling approach recently suggested that UGT71C5 is activated by persulfidation of a cysteine residue at the active site, which would add an additional post-translational layer to sulfur signaling in pathogen response (M. Li et al., 2024). The strongly cysteine-responsive UGTs of the 73B subfamily also seem to have a positive regulatory role in immune signaling since *Arabidopsis* mutant plants lacking functional UGT73B3 or UGT73B5 showed increased susceptibility to *Pst avrRpm1* (Langlois-Meurinne et al., 2005). However, the relevant substrate(s) and the mechanism behind this effect have not been identified yet.

Among the group of proteins accumulating in response to cysteine treatment as well as during pathogen infection were also several glutathione-S-transferases (GSTs) of the plant-specific phi (GSTF) and tau (GSTU) classes. GSTs catalyze the conjugation of glutathione to various substrates, mostly for detoxification purposes, and several of them are induced by pathogen infection and SA (Cummins et al., 2011; Lieberherr et al., 2003; Sappl et al., 2004). An accumulation during pathogen response has also been found in previous proteome studies (Jones et al., 2006; Maldonado-Alconada et al., 2011). However, very limited information is available on endogenous substrates or the exact metabolic functions of disease-induced GST isoenzymes. GSTF2, GSTF8, GSTF10 and GSTF11 were shown to bind to SA, though the biological implications remain unclear (Tian et al., 2012). Also, GSTF2 interacts with camalexin and was suggested to translocate plant defense compounds (Dixon et al., 2011).

Alternative oxidase (AOX) is induced by *Pst* infection and has been proposed to modulate reactive oxygen species (ROS) signaling in the mitochondria (Colombatti et al., 2014; Maxwell et al., 1999; Simons et al., 1999). The balance between AOX and manganese superoxide dismutase activities seems to be relevant for the specificity of ROS signaling by either confining the signal to the mitochondrial matrix (O_2_^−^) or spreading it to the rest of the cell (H_2_O_2_) (Cvetkoska & Vanlerberghe, 2012). Our results indicate that induction of AOX can be triggered in response to cysteine accumulation prior to or independently of a pathogen-induced ROS burst.

The general response of the *Arabidopsis* leaf proteome to pathogen attack revealed that microbe interaction induces profound changes in organelle metabolism. Mitochondria strongly increase their oxidative phosphorylation capacity and use amino acids as alternative respiratory substrates. In contrast, chloroplasts decrease most of their functions including photosynthesis as well as amino acid synthesis and focus on the production of precursors for specialized metabolites and immune signals. This general metabolic shift from growth to defense is also clearly visible in the proteome response of cysteine treated seedlings, which in addition had a slightly decreased chlorophyll content compared to the control. However, the macromolecular composition and respiration rate of the seedlings as well as the photosynthetic performance of the plants was unaffected 24h after cysteine treatment indicating that the systemic induction of defense pathways was able to protect the plant without causing major growth restrictions at this stage.

Interestingly, the protein most decreased in response to cysteine was HEM1, which has recently been reported to act as a translational regulator involved in the attenuation of the immune response during effector-triggered immunity (Y. Zhou et al., 2023). The loss of HEM1 caused exaggerated cell death and restricted bacterial growth. Thus, a major aspect of cysteine signaling during pathogen response might be the repression of antagonistic signals.

### Compartment specific functions of cysteine synthesis in immune signaling

While most amino acid anabolic pathways are localized in the chloroplasts, cysteine can be synthesized in different subcellular compartments (Heeg et al., 2008; Watanabe et al., 2008). The total leaf cysteine content after pathogen attack was reduced only in *oastl-A* mutant plants defective in cytosolic cysteine synthesis, but not in *oastl-B* or *oastl-C* indicating that the remaining isoforms can at least partially compensate for the defect. These finding confirm the quantitatively dominant role of the cytosol in cysteine production reported before (Heeg et al., 2008; Krüger et al., 2009). A previous study with a focus on cytosolic cysteine homeostasis had already demonstrated an increased susceptibility of *oastl-A* plants to *Botrytis cinerea* and *Pst* in contrast to a constitutive systemic immune response with high resistance to biotrophic and necrotrophic pathogens in *des1* mutants, which accumulate cysteine due to defects in the cytosolic degradation pathway (Álvarez et al., 2012). Our results are in good agreement with these finding. In addition, they clearly show that cysteine synthesis in the mitochondria is also crucial for *Arabidopsis* pathogen resistance. Lack of mitochondrial OASTL-C, despite constituting only about 5% of the total OASTL leaf content, had the strongest impact on pathogen response as indicated by the highest bacterial colony counts and leaf chlorosis after infection with *Pst*. Mitochondria require cysteine as a precursor for the synthesis of iron-sulfur clusters and the cofactors biotin and lipoate as well as for cyanide detoxification. It was previously reported that overexpression of the mitochondrial cysteine desulfurase NFS1, which provides inorganic sulfur for FeS-cluster synthesis, results in constitutive upregulation of defense-related genes and increased resistance against *Pst* (Fonseca et al., 2020). The mechanism behind this response is currently unknown and could be related to regulatory functions of FeS proteins. However, increased cysteine desulfurase activity would also potentially lead to an increase in the product sulfane sulfur, which, if produced in excess, can be reduced to hydrogen sulfide in the presence of a reductant like GSH (Frazzon et al., 2007). Hydrogen sulfide production by the mitochondrial β-cyanoalanine synthase CAS-C1 in turn has recently been shown to be crucial for stomatal closure in response to the pathogen-associated molecular pattern flagellin (Pantaleno et al., 2025). NFS1 was significantly induced by cysteine treatment and during *Pst* infection. Thus, an increase in compartment specific cysteine production as well as additional downstream metabolic processes seem to be involved in cysteine signaling during pathogen response.

Sulfate metabolism in plants branches at the level of adenosine 5’-phosphosulfate (APS), which can either be reduced to sulfide and incorporated into cysteine or phosphorylated by APS kinase (APK) and used for sulfation reactions. Blocking the sulfatation branch leads to an increased flux through the reductive branch of the pathway resulting in 4-fold increased leaf cysteine contents in *apk1xapk2* double knockout mutant plants compared to the wild type (Mugford et al., 2009). Re-evaluation of published datasets reveals that the expression profile of the *apk1xapk2* mutant strongly overlaps with the characteristic transcriptional response of *Arabidopsis* plants to bacterial pathogens, which would be in line with cysteine acting as a metabolic signal (Suppl. Figure S4). Since the cytosolic and mitochondrial but not the plastidic isoform of OASTL were induced in the APK deficient plants, compartment specific effects might again be relevant.

### Potential mechanisms of cysteine immune signaling

Signaling events can be mediated by receptor proteins coupled to either kinase cascades or ion channels, allosteric protein regulation or post-translational modifications. Previous proteomic studies have already demonstrated an effect of *Pseudomonas syringae* infection on post-translational modifications such as S-nitrosylation in *Arabidopsis* leaves (Jones et al., 2006; Maldonado-Alconada et al., 2011). However, the mechanism of cysteine sensing and signaling during pathogen response remains poorly understood. Emerging evidence suggests possible modes of action that warrant further investigation. Glutamate receptor-like calcium channels (GLRs) are involved in long-distance plant defense signaling in response to wounding (Toyota et al., 2018). GLR3.3 is activated by GSH and several amino acids and responds to L-cysteine in physiologically relevant micromolar concentrations (Alfieri et al., 2020; Grenzi et al., 2023). Treatment of the leaves with GSH or cysteine suppressed *Pst* propagation in wild type but not *glr3-3* mutant plants indicating that induction of pathogen response in reaction to extracellular cysteine is associated with this signaling pathway (F. Li et al., 2013). Based on a detailed transcriptome study and subsequent experimental confirmation clade 2 GLRs have also been linked to immune signaling (Bjornson et al., 2021). Since GLRs are mostly localized in the plasma membrane, they can detect extracellular changes in cysteine and other amino acids due to cell damage or release by pathogens but are unlikely to be involved in intracellular signaling. The induction of proteins associated with a DAMP-response in the cysteine treated seedlings indicates that extracellular receptors might be relevant.

Persulfidation of protein cysteine residues has been established as a post-translational modification involved in ABA signaling during drought response (Chen et al., 2020; Shen et al., 2020). A function in plant microbe interactions has not been demonstrated yet. However, in *Aspergillus fumigatus*, a human fungal pathogen causing severe pulmonary infections, persulfidation levels have been linked to both, the pathogenic potential of the fungus as well as the antifungal potency of alveolar macrophages and epithelial cells of the host (Sueiro-Olivares et al., 2021). Although the mechanisms of protein persulfidation and de-persulfidation is not entirely clear yet, cysteine most likely serves as the sulfur donor either via desulfhydration producing hydrogen sulfide or via transamination to 3-mercaptopyruvate and subsequent transsulfuration (Pedre et al., 2023; Shen et al., 2020). Thus, an increase in cysteine content might be translated to persulfidation signals. The cysteine desulfhydrase DES1 (AT5G28030), which has been previously associated with protein persulfidation in *Arabidopsis* (Aroca et al., 2017; Shen et al., 2020; M. Zhou et al., 2021), is not included in our proteomics dataset and 3-mercaptopyruvate sulfurtransferase (AT1G79230), the *Arabidopsis* ortholog of a yeast protein persulfidase (Pedre et al., 2023), was slightly decreased during *Pst* infection (Suppl. Dataset S2). Thus, there is currently no clear evidence for a role of protein persulfidation in cysteine signaling and this aspect requires further investigation. Another potential mechanism of cysteine signaling would be via protein cysteinylation, which has been demonstrated in animals (Martí-Andrés et al., 2024). Downstream effects can include transcriptional or post-translational regulation. According to our results, the modulation of phytohormones and defense compounds as well as the alteration of immune gene translation pose potential hubs for cysteine-induced biotic stress signaling.

In conclusion, cysteine, an important precursor for defense compounds, was shown to elicit plant immune signaling and thus serve as an infochemical during plant microbe interactions. Compartmental cysteine synthesis appears to be vital for properly mounting a pathogen response. The specific signaling pathways still need to be identified including potential intracellular cysteine receptors as well as connections to hormonal crosstalk during stress response. A potential role in hypoxic signaling indicated by proteomics also deserves further investigation.

## Materials and Methods

### Plant Material and growth conditions

All experiments were performed with *Arabidopsis thaliana* ecotype Col-0 as wild type control. All mutants are T-DNA insertion lines deficient (knockout) of the respective genes and were derived from Col-0. Homozygous seeds of *oastl-A* (At4g14880; N572213; SALK_072213; aka *oastl-a1.1*; characterized in López-Martín et al., 2008) and *oastl-B* (At2g43750; N521183; SALK_021183; characterized in Heeg et al., 2008) as well as of *oastl-C* (At3g59760, N500860; SALK_000860; characterized in Heeg et al., 2008) were kindly provided by Stanislav Kopriva and Andreas Meyer, respectively. Stratified seeds were sown in pots containing soil and grown in a climate chamber under short-day conditions with 8h of light (120 µmol · m^-2^ · s^-1^). The temperature during the day and night changed from 22 to 18 °C. Soil-grown plants were kept under these conditions for 6 weeks until experiments were performed.

For seedling cultures, seeds were incubated in 100% (v/v) ethanol for 2 min followed by two consecutive incubation steps in 6% (v/v) sodium hypochlorite (Carl Roth, Germany) for 2 min each. Afterwards, seeds were washed five times using sterile ddH_2_O and transferred to liquid growth media containing 0.43% (w/v) Murashige & Skoog salt mixture (Merck, Germany), 3 mM MES; and 0.5% (w/v) sucrose at pH 5.8. Individual cultures consisting of approximately 3 mg seeds in 50 ml liquid growth media were cultivated under long day conditions with 16 h of light (120 µmol · m^-2^ · s^-1^) shaking at 100 rpm. The temperature during the day and night changed from 22 to 18 °C. Seedlings cultures were kept under these conditions for 6 days until treatment.

### Cysteine treatment

For seedling cultures, 1 mM L-cysteine (Merck, Germany) was supplemented after 6 days of growth. The cysteine content in the medium decreased to low µM concentrations within the 24h treatment.

Conditions for cysteine feeding of plants grown on soil were optimized in preliminary experiments to achieve a similar increase in tissue cysteine levels as in the experiments performed with seedling culture. Pots were soaked in 1 mM MES pH 5.8 with or without 10 mM L-cysteine (Merck, Germany) for 24 hours starting at the beginning of the light period. Afterwards, plants were either harvested and flash frozen in liquid nitrogen immediately or follow-up experiments were performed.

### Quantification of total lipids

The total lipid content was determined using the sulpho-phospho-vanillin method described in (Park et al., 2016). Absorbance was measured at 530 nm using a spectrophotometer (Multiscan Sykhigh, Thermo Fisher Scientific, Germany).

### Quantification of total carbohydrates

The total carbohydrate content was determined using the phenol-sulphuric acid method described by. (Tamboli et al., 2020). 5 mg of lyophilised plant powder was dissolved in 1 ml of 2.5 N HCl and incubated for 3 h at 95 °C, shaking. The extracts were diluted (1:50) with demineralized water and 10 µl phenol and 1 ml concentrated sulphuric acid were added. After incubation (10 min, 95 °C, shaking) the absorbance was measured at 490 nm with a spectrophotometer (Multiskan Skyhigh, Thermo Fisher Scientific, Germany).

### Extraction and quantification of total protein

5 mg lyophilized seedling powder was dissolved in 700 µl methanol (100% (v/v)) and incubated for 20 min shaking at 80 °C. After centrifugation (10 min, 4 °C, 18,800 xg) the pellet was washed twice in 1 ml ethanol (70% (v/v)) and resuspended in 400 µl NaOH (0.1 M). The solution was incubated for 1h shaking at 95 °C and centrifuged again. The protein content of the supernatant was quantified using Ready-to-use Coomassie Blue G-250 Protein Assay Reagent (Thermo Fisher Scientific, Germany) and Albumin Standard 23209 (Thermo Fisher Scientific, Germany).

### Quantification of chlorophyll

The quantification of chlorophyll was carried out according to a modified version of the method described by (Lichtenthaler, 1987). 5 mg plant powder was dissolved in 700 µl methanol (100% (v/v)) and incubated for 20 min at 80 °C with shaking. After centrifugation (10 min, 4 °C, 18,800 xg), the chlorophyll content of the supernatant was quantified with a spectrophotometer (Multiskan Skyhigh, Thermo Fisher Scientific, Germany) (wavelengths: 470 nm, 653 nm and 666 nm).

### Analysis of seedling respiration

Respiration rates were measured at ambient conditions using a Clark type oxygen electrode (model DW1, Hansatech Instruments Ltd, United Kingdom). Measurements were performed on control seedlings and seedlings fed 1 mM L-cysteine for 24 h in fresh, air-saturated liquid growth medium under constant stirring. After equilibration, 1 mM L-cysteine (Merck, Germany) was added. After respiration measurements, seedlings were lyophilized and oxygen consumption rates were calculated on the basis of dry weight.

### Reactive oxygen species staining

Seedlings were rinsed with distilled water and transferred to the staining solution. For 3,3’-diaminobenzidine (DAB; Merck, Germany) staining, seedlings were incubated in DAB staining solution (50 mM sodium phosphate buffer pH 7.5; DAB 0.1% (w/v); Tween 20 0.05% (v/v)) for 7 h at 10 rpm. Afterwards, seedlings were rinsed with distilled water and incubated in destaining solution (ethanol 60% (v/v), glycerol 20% (v/v), acetic acid 20% (v/v) for 15 min at 95 °C to remove chlorophyll for proper visualization of the stain. Stained seedlings were scanned using the Epson Perfection V850 Pro (Epson, Germany).

### Glutathione-S-transferase activity assay

Glutathione-S-transferase (GST) activity in crude extracts of seedlings was determined using a 1-chloro-2,4-dinitrobenzene (CDNB)-based photometric assay and performed as stated in Koschmieder et al. (2022) if not mentioned otherwise. In short, 150 µl of cold extraction buffer (50 mM Tris-HCl, pH 8, 50 mM NaCl, 1 mM EDTA and 1% (w/v) PVPP) was added to 50 mg of seedlings ground in liquid nitrogen. Samples were resuspended thoroughly, kept on ice for 5 min and centrifuged at 18,000 xg for 15 min at 4 °C to obtain crude extracts. Protein concentration was determined using Bradford assay and 10-15 µg of enzyme was added to 200 µl GST assay (2 mM GSH, 1 mM CDNB in modified PBS). The increase in absorbance at 340 nm was monitored every 5 s for 10 min at room temperature and GST activity was calculated.

### Analysis of photosynthetic performance

For Chl a fluorescence analyses, the HEXAGON IMAGING-PAM (Walz, Effeltrich, Germany) was used. Saturation light pulses of 0.5 s were applied after 30 min dark treatment to determine Fm and Fm’ during the illumination with increasing actinic light intensities. Non-photochemical quenching (NPQ) was calculated as (Fm − Fm′)/Fm′ and Φ_PSII_ as (Fm′ − F’)/Fm (Baker, 2008).

### Pseudomonas syringae infection assays

For bacterial growth curves, *Pseudomonas syringea pv. tomato* DC3000 (*Pst*), grown on Kings-B Agar (KB) plates containing 50 µg · ml^-1^ Rifampicin, were resuspended in 10 mM MgCl_2_ to a final concentration of 5 × 10^5^ colony forming units (cfu)·ml^-1^ (OD_600_ 0.001). For shot-gun proteomics analysis, 2.5 × 10^6^ cfu · ml^-1^ (OD_600_ 0.005) *Pst* DC3000 was prepared as described. Five to seven of the youngest, fully matured leaves per plant were hand-infiltrated with the bacterial suspension using a needless syringe. For bacterial growth curves, leaf discs were cut from infected leaves 0 days past infection (dpi) and 3 dpi and ground in 200 µl sterile MilliQ water using a tissue lyser (Mill Retsch MM400, Retsch GmbH, Haan, Germany). Dilution series were plated on KB plates containing 50 µg · ml^-1^ Rifampicin and 50 µg · ml^-1^ Cycloheximide, and colony forming units were counted after two days of growth at 28 °C. Students t-tests were performed with 8 biological replicates per time point and genotype or treatment from two independent experiments showing similar results. For thiol quantification and shot-gun proteomics, all infected leaves were harvested, flash frozen in liquid nitrogen and ground for further procedures.

### Quantification of sulfur metabolites

Sulfur compounds in plant samples were derivatized using bromobimane and quantified by reverse phase high performance liquid chromatography (HPLC). Samples were solved in derivatisation buffer (1.5 mM bromobimane (Merck, Darmstadt, Germany); 32% (v/v) acetonitrile; 10.3 mM EDTA and 103 mM HEPES pH 8) and incubated at 1400 rpm for 30 min in darkness. Afterwards, 15.9 mM methanesulfonic acid was added, cell debris pelleted at 18,000 xg for 5 min and the supernatant filtered using Corning Costar Spin-X (0.22 µm) centrifuge filter tubes (Merck, Darmstadt, Germany). Samples were diluted and measured using an Agilent 1260 Infinity II HPLC (Agilent Technologies, Waldbronn, Germany) by fluorescence detection (ex. 380 nm; em. 480 nm). Peaks were evaluated and quantified using OpenLabCDS software (Agilent, Santa Clara, United States).

### Quantification of camalexin

Camalexin was determined by HPLC as described in (Koprivova et al., 2019). In short, approximately 50 mg plant material were extracted in 150 µl of dimethlysulfoxide (DMSO) for 20 min with shaking and centrifuged. 20 µl of extracts were injected into a HPLC system with a Spherisorb ODS-2 column (250 mm x 4.6 mm, 5 µm) and resolved using a gradient of acetonitrile in 0.01% (v/v) formic acid. Camalexin was detected by a FLD detector with an excitation at 318 nm and emission at 368 nm as described in (Bednarek et al., 2011). For the quantification of camalexin external standards were used ranging from 1 pg to 1 ng per µl.

### Protein extraction, digestion and sample preparation for proteome analysis via mass spectrometry

Protein extraction, purification and digestion was performed with an adapted single-pot solid-phase-enhanced sample preparations (SP3) protocol from (Mikulášek et al., 2021) which originates from (Hughes et al., 2019). In short, 5 mg of lyophilized plant powder were solved in 500 µl SDT buffer (4% sodium dodecyl sulfate, 0.1 M dithiothreitol, 0.1 M Tris pH 7.6) and incubated at 60 °C for 30 min. Thirty microliters of the supernatants were mixed with 7.5 µl iodoacetamide (0.1 M) and incubated for 30 min in the dark. Then 2 µl dithiothreitol (0.1 M) was added. Equal shares (v/v) of hydrophobic and hydrophilic carboxylate-modified magnetic beads (Sera-Mag: No. 441521050250, No. 241521050250, GE Healthcare) were prepared for protein binding. Here, 600 µg beads were used per sample. Washing steps with ethanol (80%) were performed on a magnetic rack as described in (Mikulášek et al., 2021). The bound proteins were digested for 18 h at 37 °C with 0.5 µg trypsin (MS grade, Promega) per sample. The peptide-containing supernatants were collected in tubes with low peptide binding properties. The beads were rinsed in 60 µl ammonium bicarbonate (50 mM) to recover the remaining peptides. The eluates were acidified with formic acid and desalted on 50 mg Sep-Pak tC18 columns (Waters). Peptide concentrations were quantified using the Pierce Quantitative Colorimetric Peptide Assay Kit (Thermo Fisher Scientific) and adjusted to the same concentration in 0.1% formic acid.

### Shotgun proteomics by ion mobility mass spectrometry (IMS-MS/MS)

For proteome profiling of the cysteine treated seedlings, 200 ng of peptides were injected via a nanoElute 1 (Bruker Daltonic, Bremen, Germany), separated on an analytical reversed-phase C18 column (Aurora Ultimate 25 cm×75 µm, 1.6 µm, 120 Å; IonOpticks) and analyzed with a timsTOF Pro 2 mass spectrometer. Using a multi-staged linear gradient (Eluent A: MS grade water containing 0.1% formic acid, Eluent B: acetonitrile containing 0.1% formic acid, gradient: 0 min, 2%; 54 min, 25%; 60 min, 37%; 62 min, 95%; 70 min, 95% eluent B), peptides were eluted and ionized by a CaptiveSpray 1 ion source with a flow of 300 nl min^−1^. The ionized peptides were analyzed with a standard data-dependent acquisition parallel accumulation– serial fragmentation application method (DDA-PASEF) with the following settings: Ion mobility window: 0.6–1.6 V·s/cm², 10 PASEF ramps, target intensity of 20,000 (threshold 2,500), and a cycle time of ∼1.1 s.

For proteome analysis of plants infected with *Pseudomonas syringae* 400 ng of peptides were injected via a nanoElute 2 UHPLC (Bruker Daltonic) and separated on the same type of analytical column with the same multi-staged gradient as above. Here, the eluting peptides were ionized by a CaptiveSpray 2 source and analyzed with a timsTOF-HT mass spectrometer. It was programmed with the following DDA-PASEF method parameters: Ion mobility window of 0.7–1.5 V·s/cm², 4 PASEF ramps, target intensity 14,500 (threshold 1,200), and a cycle time of ∼0.53 s.

### Data Processing and Evaluation

The ion mobility spectrometry (IMS–MS/MS) spectra from both experiments were analyzed with MaxQuant software (Cox & Mann, 2008) using default search parameters and TAIR10 (Arabidopsis.org) as database for protein identification. The calculation of label-free quantification (LFQ) values and intensity-based absolute quantification (iBAQ) values were both enabled. Data evaluation was performed using Perseus software (Tyanova et al., 2016). Proteins were excluded from further analysis if they were not detected in at least n−1 biological replicates in at least one of the sample groups. Subsequently, missing values were replaced with randomly chosen low values from a normal distribution. Significant changes were calculated from the LFQ values using Student’s t-tests (p = 0.05). Fisher exact tests were performed in Perseus to identify significantly enriched or depleted metabolic pathways as well as environmental response patterns. The metabolic pathway categories were based on a modified version of MapMan (see Suppl. Datasets S1 and S2; mapman.gabipd.org). Several RNA-seq datasets were used to annotate proteins that showed a response to different stimuli, including fungal PAMPs, DAMPs, bacterial PAMPs (Bjornson et al., 2021), hypoxia (Klecker et al., 2014), ABA treatment, JA treatment (Goda et al., 2008), BTH treatment (Wang et al., 2006), or systemic acquired resistance (Gruner et al., 2013). Transcriptomics datasets for additional abiotic stress conditions (heat, cold, drought, salt, darkness) were used as summarized in (Hildebrandt, 2018).

## Data availability

The mass spectrometry proteomics data have been deposited to the ProteomeXchange Consortium (http://proteomecentral.proteomexchange.org) via the PRIDE partner repository (Perez-Riverol et al., 2022) with the dataset identifier PXD054723.

## Supplementary Data

Supplementary Figure S1: Seedling culture - dynamics and effects of cysteine treatment. Seedling phenotype, cysteine depletion in media, oxygen consumption rate and content.

Supplementary Figure S2: Overlap between the *Arabidopsis* proteome responses to pathogen interaction and cysteine treatment.

Supplementary Figure S3: *Arabidopsis* response to cysteine treatment: Chlorophyll content, photosynthetic performance, protein, carbohydrate and lipid content, GST abundance and activity and H_2_O_2_ staining.

Supplementary Figure S4: Overlap between proteome and transcriptome responses to bacterial pathogens and cysteine accumulation.

Supplementary Dataset S1: Proteomics dataset of cysteine-treated *Arabidopsis* seedlings

Supplementary Dataset S2: Proteomics dataset of *Pst*-infected *Arabidopsis* leaves

Supplementary Dataset S3: Comparison of proteomics datasets of cysteine-treated Arabidopsis seedlings and *Pst*-infected *Arabidopsis* leaves

Supplementary Dataset S4: Raw data for Fig. 1, Fig. 2, Fig. 3 and Fig. 4.

## Acknowledgments

We thank Christina Mack, Cosima Sies, and Dagmar Lewejohann for skillful technical assistance and Isabella Rotthäuser for thorough help with plant cultivation.

## Author contributions

TMH, JM, and BH designed the research; BH, CA, and JM performed and evaluated the shotgun proteomics experiments; AK and SK quantified camalexin; UA measured photosynthetic parameters, CA analyzed seedling respiration, JM performed all other experiments, TMH, JM and BH analyzed the data; JM and TMH wrote the paper with support from all other authors; TMH agrees to serve as the author responsible for contact and ensures communication.

## Conflict of interest

The authors have no conflicts of interest to declare.

**Abbreviations**
ABA: abscisic acid
AOX: alternative oxidase
APK: adenosine 5’-phosphosulfate kinase
APS: adenosine-5‘-phosphosulfate
BTH: benzothiadiazole
CAS-C1: cyanoalanine synthase
DAMP: damage-associated molecular pattern
DES1: L-cysteine desulfhydrase; dpi, days past infection
ETHE1: sulfur dioxygenase
GLR: Glutamate receptor-like calcium channels
GSH: reduced glutathione
GST: glutathione-S transferases; hpi, hours past infection
JA: jasmonic acid
NFS1: cysteine desulfurase
OAS: O-acetylserine
OASTL: O-acetylserine (thiol)lyase
PAMP: pathogen-associated molecular pattern;
Pst: *Pseudomonas syringae* pv tomato DC3000
ROS: reactive oxygen species
SA: salicylic acid
SAR: systemic acquired resistance
SERAT: serine acetyltransferase
STR1: 3-mercaptopyruvate sulfur transferase
UDP: uridine diphosphate
UGT: uridine diphosphate-dependent glucosyltransferase.

