## Supplementary Figures S1-4 for "Cysteine signaling in plant pathogen response"

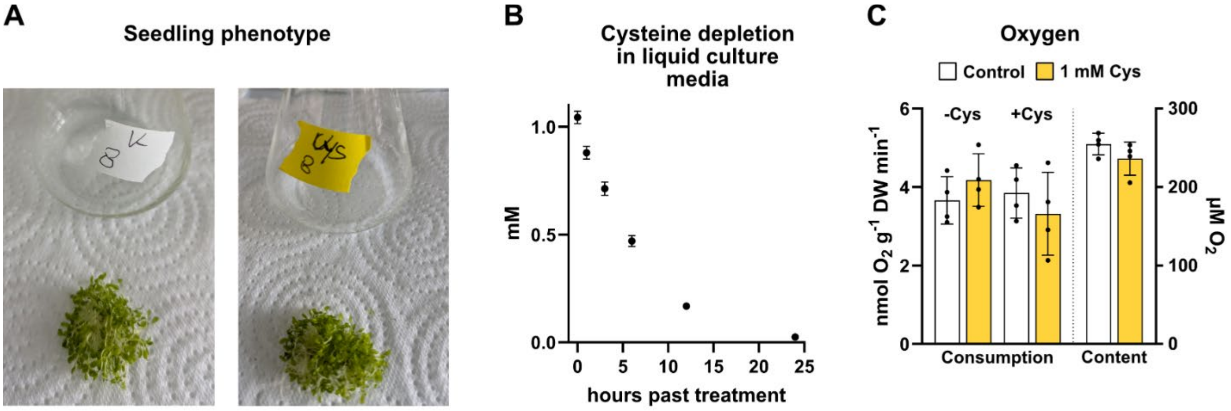


**Supplementary Figure S1: Seedling culture - dynamics and effects of cysteine treatment**.

An *Arabidopsis thaliana* seedling culture was supplemented with 1 mM L-cysteine for 24 hours before harvest. **(A)** Pictures of representative control (left) and cysteine supplemented (right) seedling cultures. **(B)** The cysteine concentration [mM] of media containing 6 days old seedlings was determined 0, 1, 3, 6, 12 and 24 hours after addition of 1 mM cysteine. Mean values (dots) ± SD are shown (n = 5). (**C**) Oxygen consumption rates [nmol $O_2$ · g dry weight$^{-1}$ · min$^{-1}$] of control seedlings (white bars) and seedlings fed with 1 mM L-cysteine for 24h (yellow bars) in the presence (+Cys) or absence (-Cys) of 1 mM L-cysteine in the respiration buffer (left axis, n = 5). The seedlings were lyophilized after respiration measurements and oxygen consumption rates were calculated based on dry weight. Oxygen content [µM] of control medium (white bar) and medium supplemented with 1 mM cysteine for 24h (yellow bar) (right axis, n = 4). Mean (bars) and individual (dots) values ± SD are shown.

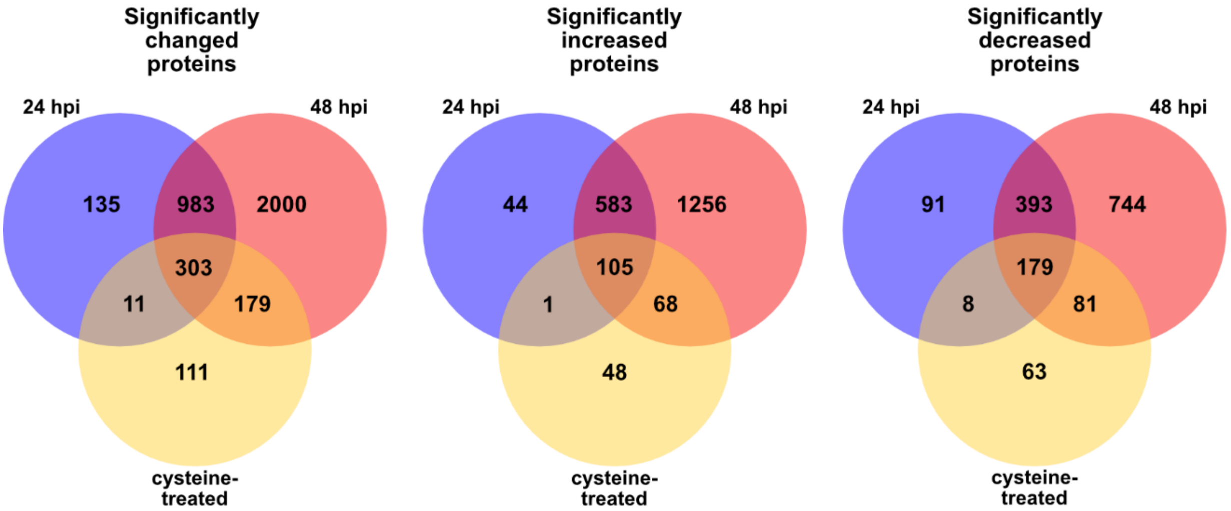


**Supplementary Figure S2: Overlap between the *Arabidopsis* proteome responses to pathogen interaction and cysteine treatment**

*A. thaliana* plants were infiltrated with *P. syringae* DC3000 (Pst) (2.5 × $10^6$ colony forming units per ml). Six leaves per plant were sampled at 24 and 48 hours after inoculation and used for shotgun proteome analysis (24 hpi and 48 hpi). In addition, an *A. thaliana* seedling culture was supplemented with 1 mM L-cysteine for 24 hours before harvest and proteome analysis (cysteine-treated). The sum of significantly increased (middle) and decreased (right) proteins in a compared subset does not necessarily equal the respective total amount of significantly changed proteins in that subset (left) due to proteins that are increased in one but decreased in the other condition or vice versa.

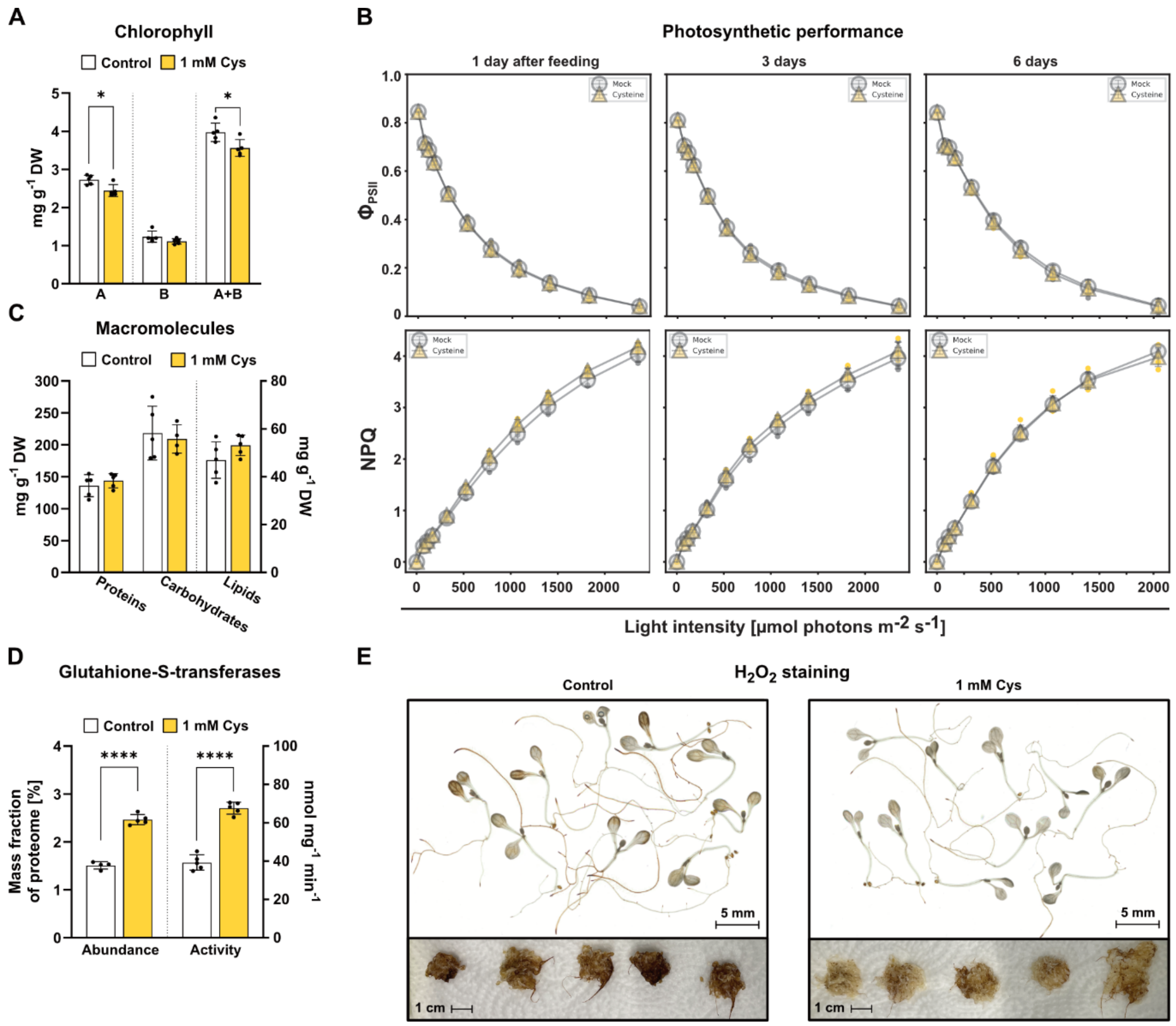


**Supplementary Figure S3:** ***Arabidopsis*** **response to cysteine treatment.**

(**A**) Seedling contents of chlorophyll A and B [mg · g dry weight-1] (n = 5). (**B**) Photosynthetic performance of rosette leaves determined via chlorophyll a fluorescence. (**C**) Seedling contents of proteins, carbohydrates, and lipids [mg · g dry weight-1] (n = 5). (**D**) Seedling abundance [mass fraction of proteome in %] and activity [nmol · $mg^{-1}$ · $min^{-1}$] of glutathione-S-transferases (n = 4-5). Mean (bars) and individual (dots) values ± SD are shown. Asterisks indicate statistically significant differences compared with control seedlings following students t-test (* P < 0.05; **** P < 0,0001). (**E**) Seedlings following $H_2O_2$ staining using 3,3'-diaminobenzidine with scans of representative seedlings (top) and pictures of whole seedling cultures (bottom).

**A,C,D,E**: An *A. thaliana* seedling culture was supplemented with 1 mM L-cysteine for 24 hours before harvest.

**B**: *A. thaliana* plants were grown for 6 weeks under short-day conditions and watered with 10 mM L-cysteine for 24 h before analysis.

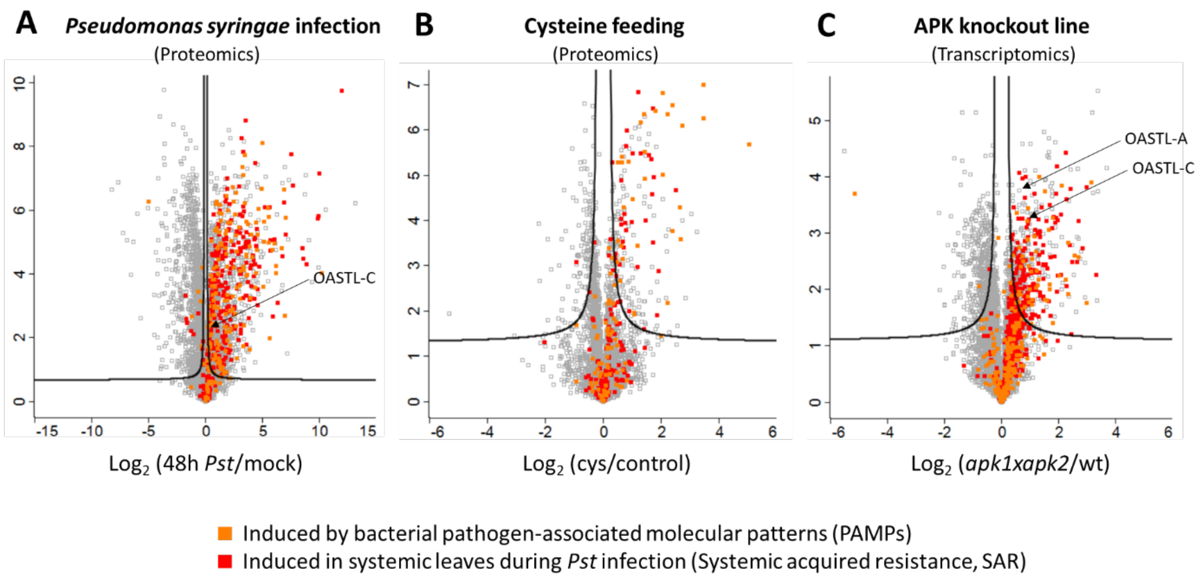


**Supplementary Figure S4: Overlap between proteome and transcriptome responses to bacterial pathogens and cysteine accumulation**

(**A**) *A. thaliana* plants were infiltrated with *P. syringae* DC3000 (Pst) ($2.5 \times 10^6$ colony forming units per ml). Six leaves per plant were sampled 48 hours after inoculation and used for shotgun proteome analysis. Infected leaves had 6.6fold increased cysteine contents compared to mock-infected leaves. **(B)** An *A. thaliana* seedling culture was supplemented with 1 mM L-cysteine for 24 hours before harvest and proteome analysis. Cysteine treated seedlings had 7.6fold increased cysteine contents compared to the control. **(C)** Transcriptome analysis of rosette leaves of adenosine 5'-phosphosulfate kinase deficient Arabidopsis mutants compared to wild type plants. Mutant plants had 4fold increased cysteine contents compared to the wild type. Data are taken from Mugford et al. (2009).

In all three volcano plots genes with significantly induced expression ($\log_2$ FC $\geq$ 1.5) in response to bacterial PAMP treatment (Bjornson et al., 2021) or during systemic acquired resistance (Gruner et al., 2013) are highlighted in orange and red, respectively.

OASTL-A: Cytosolic O-acetylserine (thiol) lyase (AT4G14880)

OASTL-C: Mitochondrial O-acetylserine (thiol) lyase (AT3G59760)
